# Differential gene expression in chicken primary B cells infected *ex vivo* with attenuated and very virulent strains of infectious bursal disease virus (IBDV)

**DOI:** 10.1101/180158

**Authors:** Katherine L. Dulwich, Efstathios S. Giotis, Alice Gray, Venugopal Nair, Michael A. Skinner, Andrew J. Broadbent

## Abstract

Infectious bursal disease virus (IBDV) belongs to the family *Birnaviridae* and is economically important to the poultry industry worldwide. IBDV infects B cells in the bursa of Fabricius (BF), causing immunosuppression and morbidity in young chickens. In addition to strains that cause classical Gumboro disease, the so-called ‘very virulent’ (vv) strain, also in circulation, causes more severe disease and increased mortality. IBDV has traditionally been controlled through the use of live attenuated vaccines, with attenuation resulting from serial passage in non-lymphoid cells. However, the factors that contribute to the vv or attenuated phenotypes are poorly understood. In order to address this, we aimed to investigate host cell-IBDV interactions using a recently described chicken primary B cell model, where chicken B cells are harvested from the BF and cultured *ex vivo* in the presence of chicken CD40L. We demonstrated that these cells could support the replication of IBDV when infected *ex vivo* in the laboratory. Furthermore, we evaluated the gene expression profiles of B cells infected with an attenuated strain (D78) and a very virulent strain (UK661) by microarray. We found that key genes involved in B cell activation and signaling (TNFSF13B, CD72 and GRAP) were down-regulated following infection relative to mock, which we speculate could contribute to IBDV-mediated immunosuppression. Moreover, cells responded to infection by expressing antiviral type I IFNs and IFN-stimulated genes, but the induction was far less pronounced upon infection with UK661, which we speculate could contribute to its virulence.

## Introduction

Given the expanding human population, set to reach 9.15 billion by 2050 (1) the global poultry industry is essential to securing enough food for the future. However, immunosuppression has a significant negative impact on the performance of birds (2) and consequently represents a key economic challenge, and a threat to global food security. While environmental stressors and poor nutrition contribute to immunosuppression, many cases in chickens are caused by infection with immunosuppressive viruses, for example infectious bursal disease virus (IBDV) (2).

IBDV is a highly contagious member of the *Birnaviridae* family that commonly ranks among the top 5 infectious problems of poultry (3). The virus causes an acute disease in chickens known as infectious bursal, or Gumboro, disease (4, 5). Infected birds typically display clinical signs such as ruffled feathers and pale comb/wattles, and they may fail to gain weight. In more severe cases, birds may lack normal social interaction and can be observed with half-shut eyes and drooping wings, as well as signs of weight loss and diarrhoea. In the most severe cases, birds may be solitary and depressed, showing no attempts to feed, and they may have a crop that is filled with water and vent feathers soiled with white faecal material. The virus has a preferred tropism for B cells, the majority of which are located in the bursa of Fabricius (BF), and infection leads to B cell depletion. As a result, birds that survive the infection are typically immunosuppressed, respond less well to vaccination, and are more susceptible to secondary infections, some of which are zoonotic and of Public Health importance. The economic impact of IBDV is therefore two-fold: firstly, due to losses associated with morbidity and mortality and secondly through indirect losses due to immunosuppression (5).

Improving the control of immunosuppressive viruses is a priority for the poultry industry and vaccination is the cornerstone to IBDV control. The use of both inactivated and live attenuated vaccines has been widespread since soon after the identification of IBDV in the 1960s. Next generation vaccines have also been licensed, based on a recombinant herpesvirus of turkey (HVT) vector, or immune complex vaccines (6). However, despite these control efforts, the infection remains endemic worldwide and new strains have emerged and spread, for example immune escape antigenic variants (7, 8) and a pathotypic variant of very virulent (vv) phenotype (9, 10), the latter causing increased severity of disease and higher mortality which can be up to 60% in some flocks, compared to 1-2% following infection with classical strains (5).

The chicken B cell is pivotal to the pathogenesis of IBDV. It is therefore crucial to characterise chicken B cell-virus interactions in order to improve our current understanding of viral pathogenesis and identify areas that can be exploited to develop novel strategies for controlling IBDV. Key questions that remain unanswered are the basis for the increased pathogenicity of the vv strain and the mechanism of attenuation of cell-culture adapted strains. However, until recently, it has not been possible to culture chicken primary B cells *ex vivo* as, when they are removed from the BF, they do not survive for long (11). Consequently, it has not been possible to perform a thorough analysis of the interactions of chicken B cells with different strains of IBDV, and many pathogenesis studies to date have been conducted *in vivo*, where birds are infected and bursal tissues are harvested at necropsy for downstream analysis of gene expression (12–17).

The gene encoding chicken CD40 ligand (chCD40L), a molecule responsible for B cell proliferation *in vivo*, was identified (18) and a soluble fusion protein containing its extracellular domain was subsequently shown to support the proliferation of chicken B cells in culture for up to three weeks (19). In 2015, Schermuly et al. showed that chCD40L-treated B cells could be infected with Marek’s disease virus (11), demonstrating that the cells have the potential for studying the consequences of lymphotropic virus infection. Despite these successes, chicken primary B cell cultures have not yet been applied to the study of IBDV. Here we report the successful culture of chicken primary B cells *ex vivo* in the presence of soluble chCD40L and provide data demonstrating that these cells can support the replication of an attenuated cell-culture adapted strain of IBDV (D78) and a very virulent strain that does not replicate in non-lymphoid cells (UK661). Furthermore, we characterise the gene expression profile of B cells infected with both strains of virus, identifying differences that correlate with pathogenicity.

## Results

### Chicken primary B cells can be cultured in the presence of chicken CD40L

Consistent with previous reports (11, 19), we found that when chicken primary B cells were cultured in the presence of soluble chCD40L, the number and viability of the cells was significantly increased compared to when cells were cultured in the absence of chCD40L (Fig. 1). The number of cells increased 4-fold from 9.02 x10^5^ to 3.63x10^6^ per ml over a period of 6 days when chCD40L was added to the culture media, in contrast to when it was absent (p <0.05) (Fig. 1(a)). Cell viability was also significantly improved, for example from 25% at day 3 post-culture in the absence of chCD40L to 48% in the presence (p <0.05) (Fig. 1(b)).

**Figure 1.**
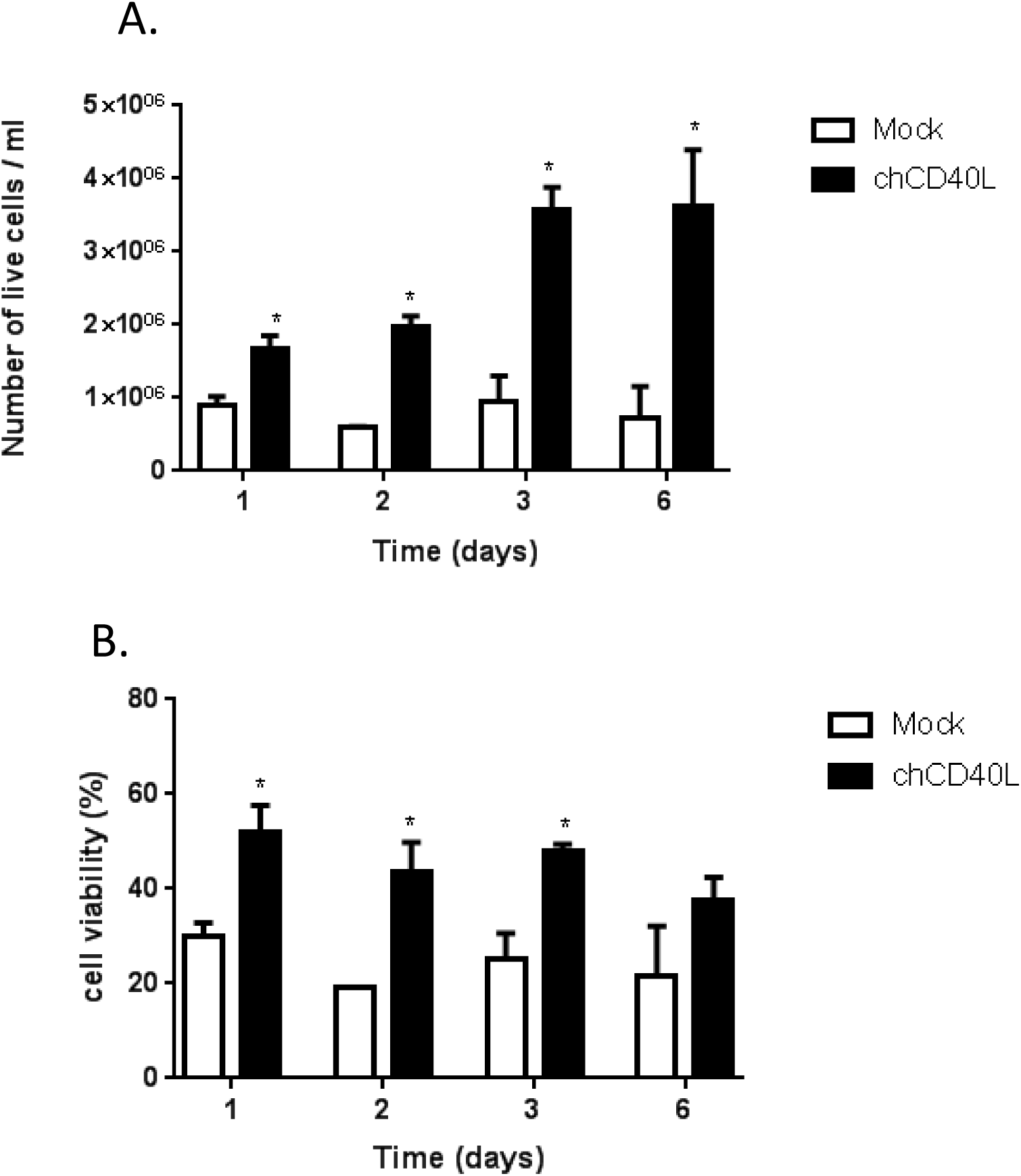
Chicken primary B cells can be cultured in the presence of chicken CD40L. Chicken primary B cells were cultured in the presence (black bars) or absence (white bars) of chicken (ch)CD40L and the number of live cells (A) and the percentage of viable cells (B) determined at the indicated time-points post infection. Data shown are representative of at least three replicate experiments, error bars represent the standard deviation of the mean, and statistical significance was determined using a paired Student’s t-test at each time point, *p<0.05.

### Chicken primary B cells can support the replication of both cell-culture adapted and very virulent strains of IBDV

At 18 hours post-infection, samples of mock and infected cell cultures were fixed, labelled with a mouse monoclonal antibody against IBDV VP2 and a goat-anti-mouse secondary antibody labelled with Alexa Fluor 488, and counterstained with DAPI. Some cells had evidence of green fluorescence around the nucleus (Fig. 2(a)), consistent with the presence of IBDV in the cytoplasm of infected cells. This was evident for both D78 and UK661 (Fig. 2(a)). RNA was extracted from infected cultures at 5, 18, 24 and 48 hours post-infection, and subjected to reverse transcription quantitative polymerase chain reaction (RTqPCR) with primers specific to a conserved region of the IBDV VP4 gene. The expression of VP4 was first normalised to a house-keeping gene (TBP) and then expressed as fold change in copy number relative to mock samples in a ΔΔCt analysis. The average fold change in IBDV VP4 expression increased to 16,603 copies at 48 hours post-infection with D78, and 38,632 copies at 48 hours post-infection with UK661. Taken together, these data demonstrate that the chicken primary B cells could support the replication of cell-culture adapted and vv IBDV strains. This is in contrast to primary chicken embryo fibroblasts (CEFs) or the immortalised chicken fibroblast cell line, DF-1, which do not support the replication of vv IBDV without prior adaptation that can lead to viral attenuation (20).

**Figure 2.**
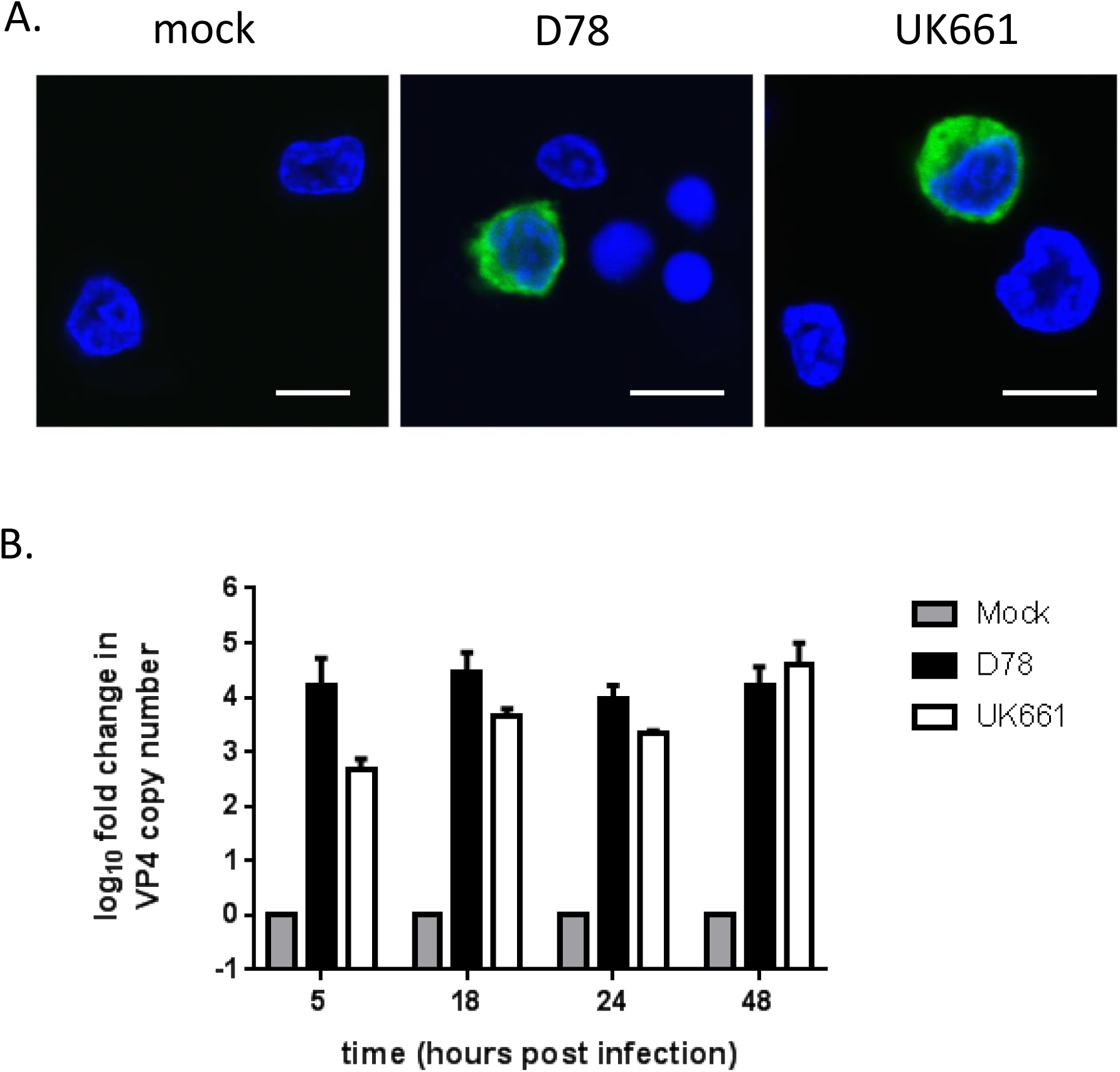
Chicken primary B cells can support the replication of both cell-culture adapted and very virulent strains of IBDV. Chicken primary B cells were mock-infected or infected with either D78 or UK661 and a sample from each culture was fixed, labelled and imaged: IBDV VP2, green; nuclei, blue; scale bar, 7μm (A). RNA was extracted at the indicated time-points post-infection, reverse transcribed and a conserved region of the IBDV VP4 gene was amplified by quantitative PCR (B). The log_10_ fold change in VP4 copy number was normalised to the TBP house-keeping gene and expressed relative to mock-infected samples as per the 2^−ΔΔCT^ method. Data shown are representative of at least three replicate experiments, and error bars represent the standard deviation of the mean.

### Chicken primary B cells infected with vv UK661 and attenuated D78 show a differential gene expression profile

Next, we aimed to evaluate how the primary B cells responded to infection with either D78 or UK661. At 5, 18, 24 and 48 hours post-infection, we determined the level of expression of type I IFN (IFNβ) and the interferon stimulated gene (ISG) IFIT5 by RTqPCR. The cells infected with D78 expressed significantly more IFNβ and IFIT5 than cells infected with UK661 at 18, 24 and 48 hours post-infection (*p<0.05, ***p<0.001, ****p <0.0001) (Fig. 3). Moreover, at these time points, the average expression of IFNβ in cells infected with UK661 was actually reduced relative to mock-infected cells, (****p<0001 at 18 hours post-infection) (Fig. 3(a)).

**Figure 3.**
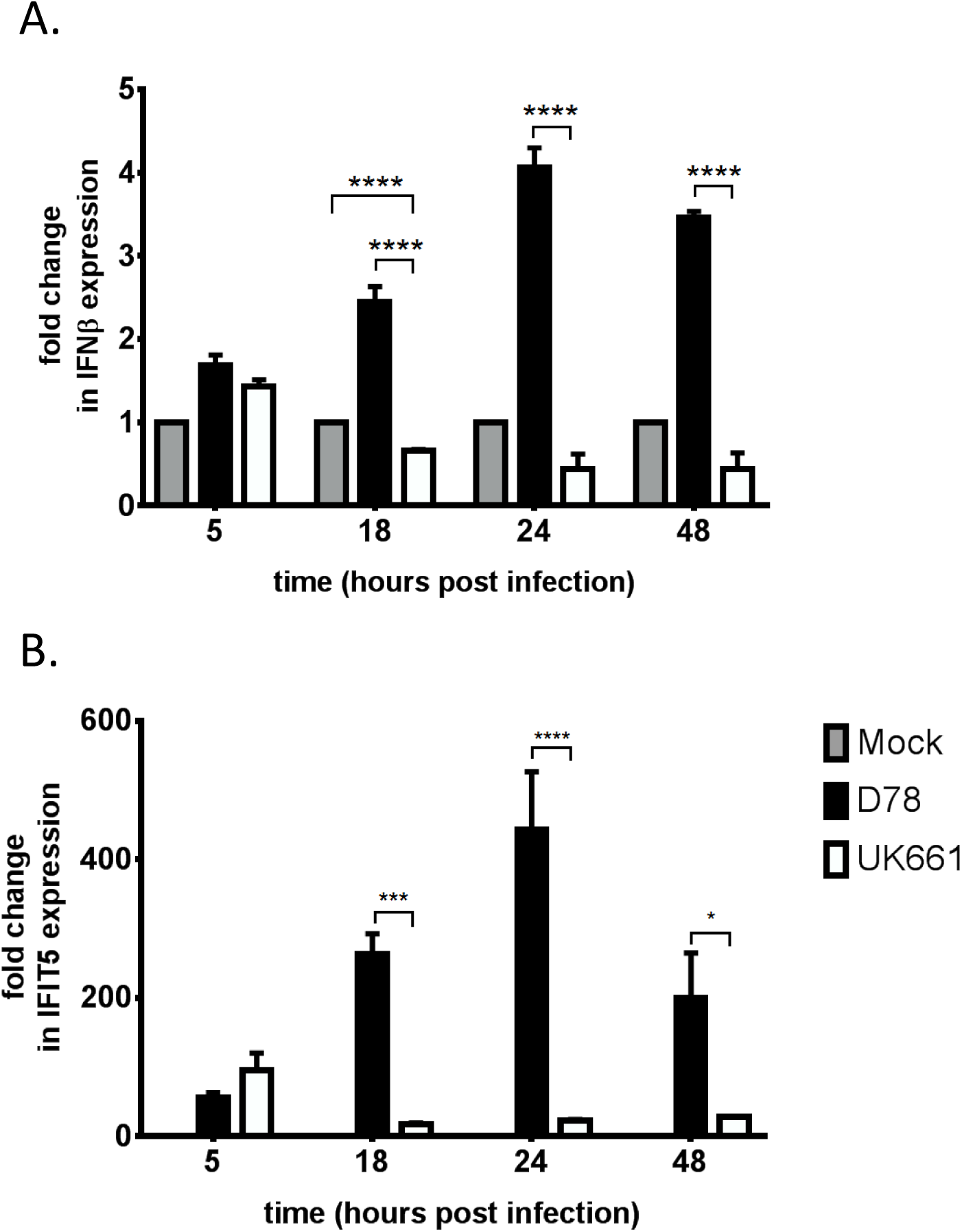
The expression of IFNβ and IFIT5 is significantly up-regulated in chicken primary B cells infected with D78 compared to cells infected with UK661. RNA was extracted from mock-, D78- and UK661-infected cultures at the indicated time points post-infection, reverse transcribed and the chicken IFNβ (A) and IFIT5 (B) genes were amplified by quantitative PCR. The fold change in the expression of each gene was normalised and expressed relative to mock-infected samples as per the 2^−ΔΔCT^ method. Error bars represent the standard deviation of the mean and statistical significance of the difference in gene expression in D78-infected cultures compared to UK661-infected cultures was determined by using a two-way ANOVA (*p<0.05, ***p<0.001, ****p <0.0001).

To acquire a broader understanding of the effect of infection on the transcriptome, gene expression of mock-, D78- and UK661-infected cultures was screened at 18 hours post-infection using the Affymetrix Chicken Genome array. This array contains comprehensive coverage of 32,773 transcripts corresponding to over 28,000 chicken genes and also includes probe sets for detecting IBDV transcripts, which we used to confirm the virus infection. Full raw and processed microarray data have been deposited in ArrayExpress with the Accession Number E-MTAB-5947. Principal-component analysis (PCA) was used to visualize three-dimensional expression patterns of the RNA data sets (Fig. S1). The samples for each individual treatment (mock-, D78-, and UK661-infected samples) mapped near to each other in a cluster, reflecting minor variations within replicates of each treatment. However, the groups were mapped separately to one another, demonstrating their transcriptomic distinctiveness from one another.

Analysis of the array data showed that 69 genes were differentially regulated, relative to mock-infected cells, following D78 infection (*p*<0.05, fold change cut-off: 1.5), 12 of which were also those differentially regulated following infection with UK661 (Fig. 4(a)). Of the 69 genes differentially regulated following D78 infection, 53 were up-regulated and 16 were down-regulated. In contrast, all 12 genes differentially regulated following UK661 infection were up-regulated; there were no statistically significant down-regulated genes.

**Figure 4.**
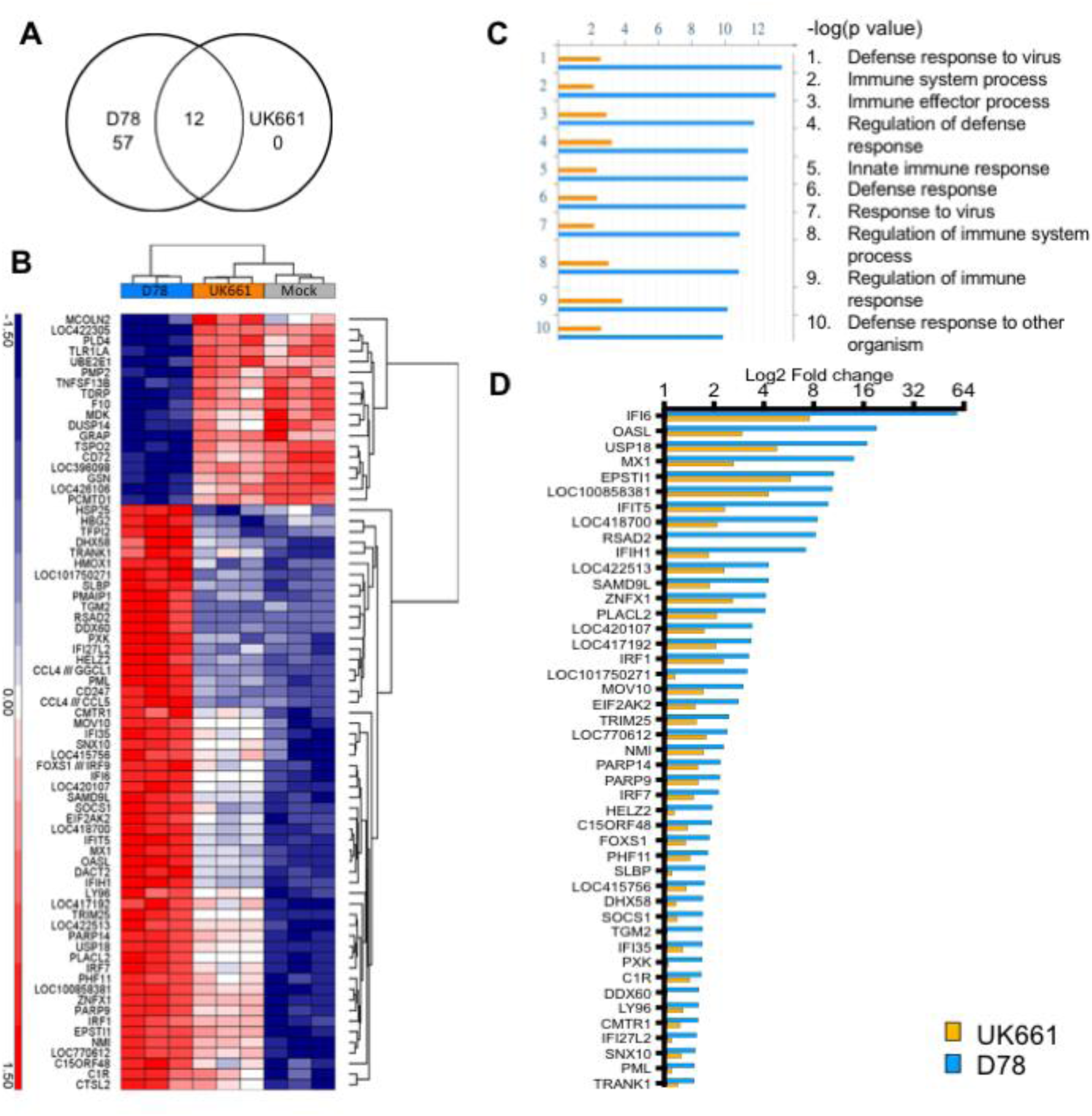
Chicken primary B cells infected with UK661 and D78 have a differential gene expression profile by microarray. RNA was extracted from mock-, D78- and UK661-infected cultures at 18 hours post-infection, and subject to microarray analysis. (A) Venn diagram showing the overlap of differentially expressed genes by D78 and UK661 relative to mock-infected cultures. (B) Hierarchical Clustering heat map of 73 transcripts that were found to be significantly regulated in at least one of the study’s microarray gene expression comparisons using Partek v6.6 software: D78 Vs Mock (blue), UK661 Vs Mock (orange) and D78 Vs UK661 (gray). Each column represents a sample, and each row represents mRNA quantification of the indicated transcript. The default settings of Euclidian dissimilarity were used for each row (log_2_ transformed and median centred transcripts). The colour corresponds to expression intensity values as indicated by the vertical legend (red, up-regulated; dark blue, down-regulated). (C) Distribution histogram of GO process term enrichment analysis using the MetaCore enrichment analysis tool. The ten most enriched GO process terms in the microarray comparisons D78 (blue) and UK661 (orange) were plotted relative to mock, as sorted by “statistically significant maps,” which list terms in decreasing order of the standard deviation of the -log (p-Value) between the two comparisons. (D) Comparison of the relative expression of known chicken ISGs between B cells infected with D78 (blue) and UK661 (orange).

A direct comparison of gene expression between D78 and UK661 infected samples (Table S4) identified 37 differentially regulated genes, 27 of which were up-regulated by D78, relative to UK661 infection, and 10 of which were down-regulated. Two of the D78 versus UK661 up-regulated genes (HBG2 and HSP25), and two of the down-regulated genes (LOC422305 and MCOLN2) were not identified by the comparisons with mock-infected cells.

Unsupervised hierarchical clustering analysis of the significantly differentially regulated genes of the study confirmed that more transcripts were up-regulated following D78-infection compared to UK661-infection (Fig. 4(b)). All the genes could be divided broadly into four similarly-sized groups: The first group included the 16 genes that were transcribed at lower levels in D78-infected samples compared to mock-infected samples. These genes were involved in B-cell activation and signalling (TNFSF13B (which encodes BAFF), CD72, GRAP), immune processes (TLR1LA, DUSP14, PLD4, MDK, PMP2, F10, GSN), and other processes such as protein ubiquitination (UBE2E1) and cholesterol transport and binding (TSPO2). In addition, LOC422305 and MCOLN2, found to be significantly down-regulated in cells infected with D78 relative to cells infected with UK661, were also included in this group, making a total of 18 genes. The LOC422305 gene encodes the mitochondrial-like ES1 protein (21), and the MCOLN2 gene encodes the ion channel TRPML2 (22) that has been reported to enhance the replication of yellow fever and dengue viruses (23, 24). The other three groups comprised the 53 genes that were transcribed to a higher level in D78-infected cells relative to mock-infected cells, as well as the two genes that were significantly up-regulated by infection with D78 relative to infection with UK661 (HBG2 and HSP25), making a total of 55 genes. These were further sub-grouped on the basis of their expression in cells infected with UK661, with their transcripts being present at: a similar level (19 genes), a marginally higher level (17 genes) or a moderately higher level (19 genes) than in mock-infected cells. Most of the genes in these groups were involved in innate immune responses (IFI6, SAMD9L, NMI, IRF7, LGP2, IFI35, HMOX1, LY96, SOCS1, and others), antiviral responses (OASL, Mx1, IFIT5, RSAD2, IFIH1, IRF1, EIF2AK2 or PKR, TRIM25, IRF7, PMAIP1, LGP2, DDX60 and PML), TGFβ signalling (DACT2, PML), inflammation (CCL4, CCL5) and apoptosis (PMAIP1, PML). Previous reports have also shown that members of several virus families modulate HSP25 expression in a variety of cell types suggesting it has a broad role (25).

Enrichment analysis was performed using MetaCore (Clarivate) by matching differentially regulated genes in functional Gene Ontologies (GO) and biological processes. The probability of a random intersection between a set of genes with ontology processes was estimated with the “P” value of the hypergeometric intersection (See Fig. S2-S4 for a more detailed analysis). The top GO processes were “defence response to virus” and “immune system processes” (Fig. 4(c)). Taken together, these data show that following IBDV infection cells launch a type I IFN response, characterised by the induction of ISGs, but that the response is more marked following D78-infection than UK661-infection. One limitation of the enrichment analysis is that it relied on a comparison with mammalian gene counterparts. Previously, we have defined chicken ISGs by using DNA microarrays and RNA sequencing in CEFs stimulated with recombinant chicken IFNα (26), and we have made the full list of chicken ISGs publicly available at http://cisbic.bioinformatics.ic.ac.uk/skinner. To confirm and extend the enrichment analysis observations, we compared the relative expression of known chicken ISGs between cells infected with the two virus strains. While the core set of virus-induced genes was similar, the magnitude of the change in gene expression was lower in cells infected with UK661 than in cells infected with D78. Moreover, RSAD2, TGM2 and DDX60 were not induced by UK661-infection (Fig. 4(d)).

### Confirmation of microarray data for selected genes by real-time RT-PCR

The microarray results were validated by determining the level of expression of a panel of genes (IFIT5, IFI6, OASL, Mx1, RSAD2, IFIH1, IFNα and IFNβ) by RTqPCR (Fig. 5). For this analysis, the log_10_ of the fold changes by microarray and RTqPCR were plotted against one another and the Spearman rank correlation calculated. The mean Spearman correlation coefficient (Spearman’s *ρ*) of cDNA microarray versus qPCR was 0.90. Therefore, the level of differential gene expression detected by each platform was highly similar.

**Figure 5.**
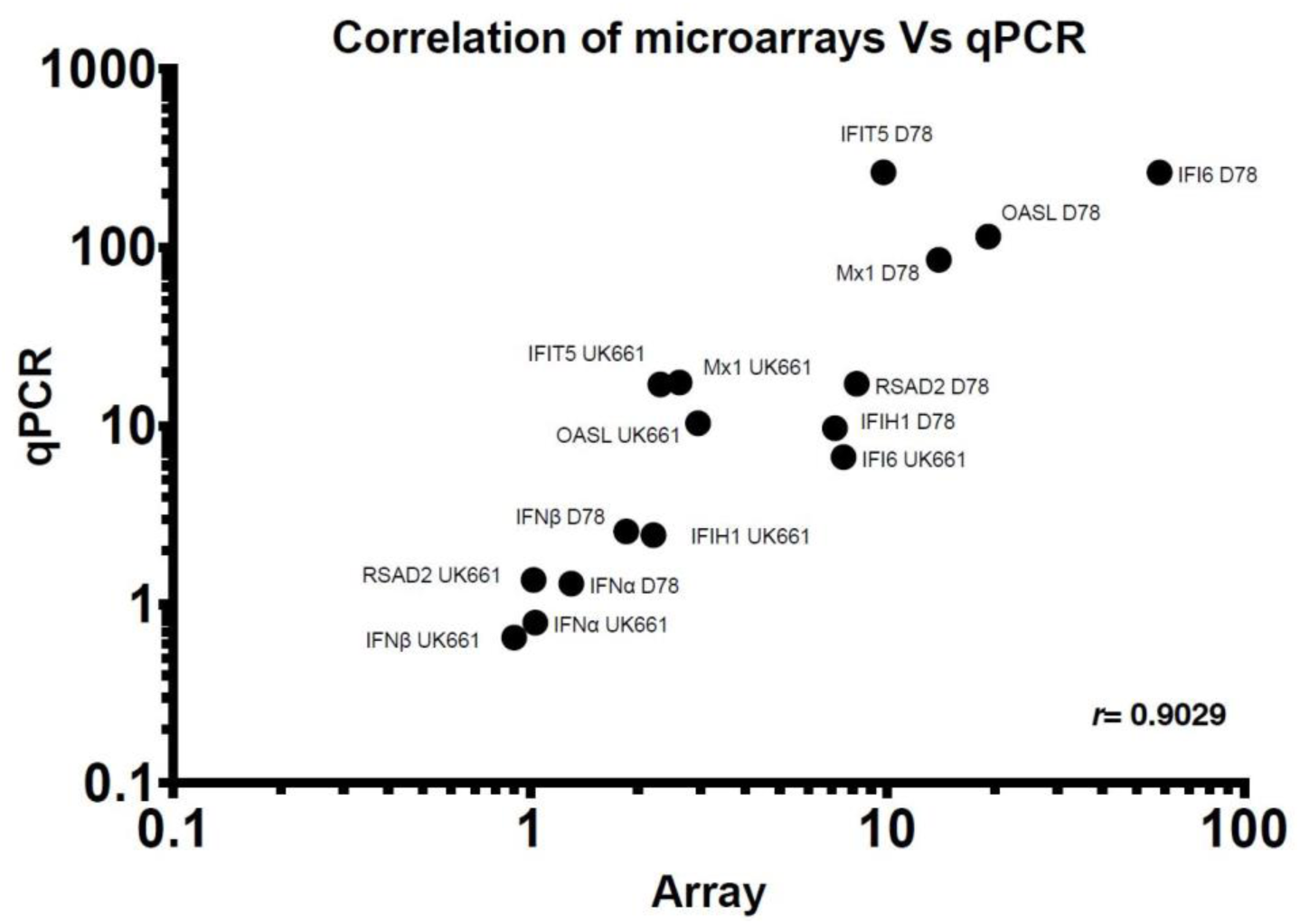
Validation of the microarray results by RTqPCR. The log_10_ fold change of a panel of innate immunity related genes (IFIT5, IFI6, OASL, Mx1, RSAD2, IFIH1, IFNα and IFNβ) was determined by microarray and RTqPCR, and plotted against one another. GraphPad Prism v7.0a was used to calculate the mean Spearman correlation coefficient (*r*).

Taken together, our data show that chicken primary B cells, cultured *ex vivo* in the presence of chCD40L, can support the replication of cell-culture adapted and vv strains of IBDV. We identified several genes involved in B cell activation and signalling that were down-regulated following D78 infection, including TNFSF13B (the gene encoding BAFF), CD72 and GRAP. The cells responded to the infection by inducing a type I IFN response with the up-regulation of ISGs, however, the magnitude of the response was considerably lower in cells infected with the vv strain UK661 than in cells infected with the attenuated strain D78.

## Discussion

Despite decades of vaccination, IBDV continues to be endemic world-wide and frequently ranks in the top 5 diseases of poultry (3). Vaccines fail to induce sterilising immunity, and vaccinated birds may continue to shed wild-type strains and reassortant viruses that contain genome segments from vaccine and field strains (27, 28). Moreover, antigenically variant viruses have emerged on several occasions, necessitating the development of novel vaccines (7, 8) and a pathotypic variant, the very virulent strain, emerged in the late 1980s (apparently by reassortment) to spread across the globe, often necessitating the use of less attenuated vaccines (6, 9, 10). Even in the absence of overt clinical signs, immunosuppression caused by IBDV infection can be an underlying cause of increased susceptibility to secondary infections, some of which can be zoonotic. For example, pre-exposure of chickens to IBDV exacerbated the pathology, persistence, or shedding of *Salmonella enteritidis, Campylobacter jejuni* and *Escherichia coli* (29–31), and a recent case-control study in Pakistan identified a history of IBDV infection in flocks as a significant risk factor associated with avian influenza virus (AIV) infection in chickens (32). Moreover, experimental inoculation with IBDV prolonged the shedding of a subsequent AIV challenge (33), and it was possible to adapt a mallard H5N2 AIV to chickens by serial passage in birds that had previously been inoculated with IBDV, but not in birds that lacked IBDV exposure (34).

Improved control strategies are therefore needed and, in the future, it may be possible to breed or engineer chickens that are resistant to IBDV disease, immunosuppression or even infection, but key scientific questions need to be answered before progress can be made in these areas. IBDV has a preferred tropism for B cells, and it is crucial to develop a greater understanding of the molecular interactions of IBDV with this cell type. The majority of B cells are located in the BF and *in vivo* studies have shown that, following IBDV infection, there is an increase in the expression of genes involved in pro-inflammatory cytokine responses, type I IFN induction and signalling as well as apoptosis in the BF (12, 16, 17). However, the BF is made up multiple cell types and, following infection, there is an influx of inflammatory cells and effector T cells into the organ. The various cell populations differ in the profile of genes they express, for example elevated expression of CD3, IL-18R1 and iNOS have been attributed to an influx of T cells, NK cells and macrophages, respectively (16). It is therefore difficult to interpret how the infected B cells respond to IBDV. To address this, some research groups have characterised the transcriptional response of cells infected with IBDV in culture (35–39). These studies have the advantage of well-defined multiplicities of infection (MOIs) and time points post-infection, and as transcriptional responses are from one cell population, they are not confounded by multiple cell-types found *in vivo*. However, until recently, it was not possible to culture chicken primary B cells *ex vivo*, so the transcriptome has typically been characterised in either fibroblast cells (35, 36, 39) or dendritic cells (37). While providing some insight into host cell-IBDV interactions, vv IBDV strains will not replicate in fibroblast cells without prior adaptation and, as infection of B cells is crucial to the pathogenesis of IBDV, the relevance of the data should not be over-interpreted. Only one study to date has characterised the transcriptional response of B cells to IBDV infection (38), however that study utilised an immortalised B cell line that had been transformed following infection with ALV (DT40 cells), limiting the conclusions that can be made.

In this study, we describe the successful culture of chicken primary B cells *ex vivo* in the presence of soluble chCD40L and demonstrate that these cells can support the replication of an attenuated strain and the prototypic vv strain of IBDV. This *ex vivo* model of IBDV infection allows us to study the interactions of the virus with its relevant host cell, retaining the advantages of *in vivo* studies without the limitations of multiple cell-types that might confound the data. Moreover, our model permits the use of defined MOIs and time-points, thus retaining the advantages of *in vitro* systems. We have extended this work by describing, for the first time, the transcriptional profile of chicken primary B cells infected with IBDV. While the validity of the model in recapitulating *in vivo* data should be evaluated experimentally, the gene expression profile we have observed is consistent with that reported in previous studies. Thus 35 of the genes that we identified as being differentially expressed in D78-infected cells compared to mock-infected cells were previously reported as differentially expressed following IBDV infection *in vivo* (12, 14–17). Only one contradiction was found: the GSN gene was found to be down-regulated in our study, yet up-regulated in one *in vivo* study (17). GSN encodes Gelsolin, which regulates actin assembly and has also been associated with inhibiting apoptosis (40). The reason for this discrepancy is unknown; it could be because our experiment characterised gene expression at 18 hours post-infection, whereas the *in vivo* study was conducted at 3 and 4 days post infection.

We demonstrated that IBDV infection led to the down-regulation of key genes involved in B cell activation and signaling, such as TNFSF13B, CD72 and GRAP. This is consistent with previous *in vivo* studies that showed down-regulation of TNFSF13B, CD72 and GRAP in bursal tissue following infection with IBDV strain F52/70 (16, 17), and with an *in vitro* study that found CD72 to be down-regulated following IBDV infection of DT40 cells (38). TNFSF13B encodes the B cell activating factor (BAFF), which is essential for the survival of B cells. BAFF is a transmembrane protein that is readily cleaved to release a soluble factor. There are three known receptors for BAFF that are expressed on B cells in varying amounts related to the maturation of the cell, and BAFF-induced signaling stimulates B cells to undergo proliferation and counters apoptosis (41). CD72 is a co-receptor on B cells, the expression of which is enhanced by BAFF (42). It contains two immunoreceptor tyrosine-based inhibitory motifs (ITIMs), ITIM1 and ITIM2, and studies have associated it with both positive and negative signaling (43). Ligation of CD72 has been shown to induce the proliferation of both resting and activated B cells, with positive signaling thought to be mediated via the Grb2-Sos-Ras pathway (43). Grb2 and the Grb2-related adaptor protein (GRAP) are both members of the same family of adapter proteins that couple signals from receptor tyrosine kinases to the Ras pathway (44). Moreover, in the chicken DT40 B-cell line, the Grb2 protein has been shown to interact with the ITIM2 motif of the avian CD72 homologue, attenuating negative signals from the ITIM1 motif, thus permitting positive signaling (45). We speculate that, as BAFF leads to the up-regulation of CD72, which positively signals via Grb-2 like proteins, the combination of BAFF, CD72 and GRAP leads to B cell activation and proliferation. As all three proteins are down-regulated following IBDV infection, it is tempting to conclude that IBDV targets this pathway, thereby reducing B cell activation and proliferation, and that this could contribute to the immunosuppression observed during infection, although this requires experimental confirmation.

We demonstrated that chicken primary B cells responded to infection with D78 by up-regulating the expression of type I IFN genes and ISGs. This is consistent with a study by Hui et al, which characterised the gene expression of DF-1 cells infected with the D78 strain (35). Twenty two of the genes that we identified as being differentially expressed in D78-infected chicken primary B cells were similarly differentially expressed following D78-infection of DF-1 cells. Notably, Hui *et al* identified that D78 triggered an IFIT5-IRF1/3-Viperin pathway in infected DF-1 cells (35). In our study, D78 infection also led to the significant up-regulation of IFIT5, IRF1, IRF7 and RSAD2 (Viperin), suggesting that this pathway is also triggered in chicken primary B cells infected with D78. In contrast, this pathway did not appear to be induced by infection with UK661.

We also found that B cells infected with the attenuated strain of IBDV (D78) up-regulated the expression of genes involved in the innate immune response to a significantly greater extent than cells infected with the vv strain (UK661). Previous work using RTqPCR also showed the expression of type I IFNα and β to be significantly lower in BF tissue taken from birds infected with UK661 than with a classical strain, F52-70 (46). However, the authors studied each virus in separate experiments and inoculated birds with different titres of the two strains, limiting the conclusions that could be made. In contrast, we characterised responses in cells infected at the same time with the same MOI of the two strains. Moreover, we found that IFNβ expression was significantly reduced in cells infected with UK661 compared to mock infected cells. This suggests that UK661 is not only able to inhibit the up-regulation of type I IFN, but might actually suppress its induction. Taken together, our results imply that UK661 is able to inhibit the up-regulation of antiviral responses to a greater extent than D78, which we speculate could contribute to its enhanced virulence. To date, two components of IBDV, VP3 and VP4, have been implicated in the suppression of innate immune responses to IBDV infection. The VP3 protein binds the viral double-stranded (ds) RNA genome and is thought to block the interaction of MDA5 with the dsRNA, thereby inhibiting downstream events that culminate in type I IFN production (47). In addition, the VP4 protein binds to the cellular glucocorticoid-induced leucine zipper (GILZ) protein, which inhibits the activation of nuclear factor kappa enhancing binding (NF-KB) protein and activator protein-1 (AP-1) (48). In mammalian cells, NF-KB and AP-1 co-operate with interferon regulatory factor (IRF) 3 and 7 to stimulate type I IFN transcription. IBDV VP4 may therefore inhibit type I IFN responses via binding to and enhancing the inhibitory action of GILZ (48). It is possible that differences in the sequences of these viral proteins between D78 and UK661 lead to differences in the antagonism of type I IFN induction.

In conclusion, the application of soluble chCD40L has opened up the possibility of studying viral pathogenesis in chicken primary B cells. While cells treated in this way have previously been used in the study of the Marek’s disease herpesvirus, this represents the first study applying them to the study of IBDV-host cell interactions. Our data illustrate interesting differences between how attenuated and very virulent strains behave in infected B cells that could shed light on both the molecular basis of IBDV-mediated immunosuppression, and on the basis of the enhanced virulence of the vv strain.

## Methods

### Immortalised cell culture and virus

DT40 cells (49) were maintained in RPMI media supplemented with L-glutamine and sodium bicarbonate (Sigma-Aldrich), 10% heat-inactivated foetal bovine serum (FBS) (Sigma-Aldrich), tryptose phosphate broth (Sigma-Aldrich), sodium pyruvate (Sigma-Aldrich), and 50mM beta-mercaptoethanol (Gibco). HEK-293T cells stably expressing a soluble chicken CD40L construct, msCD8-CD40L (18), were cultured in RPMI supplemented with L-glutamine, sodium bicarbonate, 10% heat-inactivated FBS, and 1ug/ml puromycin (Gibco). Immortalised cells were cultured at 37°C and 5% CO_2_. A cell-culture adapted attenuated strain, D78 (8), and a vv strain, UK661 (9, 10), of IBDV were kind gifts from Dr Nicolas Eterradossi, ANSES, France.

## Virus titration

Ten-fold serial dilutions of D78 and UK661 viruses were added to DT40 cells in 96 well-plates and incubated for 3 days, whereupon cultures were fixed with 4% paraformaldehyde and labelled with a mouse monoclonal antibody against IBDV VP2 (clone JF7-PD5; Kim Wark PhD Thesis, University of Hertfordshire, 2000; M. A. Skinner, unpublished) and a goat-anti-mouse monoclonal antibody conjugated to Alexa Fluor® 488 (Thermo Fisher Scientific). Cultures were scored as positive or negative based on the presence or absence of green fluorescent cells and the dilution at which 50% of the cells were infected (the tissue culture infectious dose 50, TCID_50_) was computed using the method described by Reed and Muench (50).

## CD40L stock production

Supernatant was harvested from HEK-293T cells stably expressing the msCD8-CD40L construct (chicken (ch) CD40L) and concentrated using Pierce™ Protein Concentrators PES (Thermo Fisher Scientific) according to the manufacturer’s instructions. Concentrated chCD40L was filter sterilised using a 0.22μM Millex-GP Syringe Filter (Merck), stored at 4°C and titrated prior to use. Briefly, primary B cells were cultured in the presence of two-fold serial dilutions of chCD40L for up to 6 days. Each day, the number and viability of the cells was determined by Trypan Blue (Sigma-Aldrich) exclusion and the chCD40L dilution which gave the most favourable number and viability was selected for use (1:40).

## Chickens

Embryonated eggs of the Rhode Island Red line were provided by the National Avian Research Facility (NARF). Chickens were hatched and housed in specific pathogen free (SPF) facilities at The Pirbright Institute and reared until 3 weeks of age, whereupon they were humanely culled by a schedule 1 procedure. All procedures were performed in accordance with the UK Animal (Scientific Procedures) Act 1986 under Home Office Establishment, Personal and Project licences, after the approval of the internal animal welfare and ethical review board (AWERB).

## Chicken primary B cell culture

The bursa of Fabricius was harvested aseptically post mortem and washed in sterile PBS. An enzyme solution containing 2.2mg/ml collagenase D (Sigma-Aldrich) in Hanks Balanced Salt Solution supplemented with calcium (Gibco) and 7.5% sodium carbonate was used to digest the bursal tissue. Once digested the tissue was passed through a 100μM Falcon™ cell strainer (Thermo Fisher Scientific) into cell medium containing Hanks Balanced Salt Solution (Gibco) supplemented with 7.5% sodium carbonate and 500mM EDTA (Sigma-Aldrich), before pelleting. Cells were resuspended in Iscove’s Modified Dulbecco’s Medium (IMDM) supplemented with FBS, chicken serum (Sigma-Aldrich), insulin transferrin selenium (Gibco), beta-mercaptoethanol (Gibco), and penicillin/streptomycin and nystatin (B cell media; as described by (19)) before undergoing centrifugation at 2,000rpm for 20 minutes at 4°C over Histopaque 1083 (Sigma-Aldrich). Bursal cells that banded at the interface of the medium and histopaque were collected, washed in PBS and counted using the TC20™ Automated Cell Counter (Bio-Rad). Cells were seeded in 24 well plates at 1x10^7^ cells per ml and maintained at 37°C in 5% CO_2_ in B cell media supplemented with chCD40L.

## Virus inoculation

Chicken primary B cells were infected with the attenuated strain D78 and the vv strain UK661 at a MOI of 3, after which cells were washed with media and cultured in B cell media supplemented with chCD40L. Following incubation at 37°C with 5% CO_2_ for the indicated amount of time, a 100μl sample of each well was obtained for processing for bioimaging. The remaining cells were washed in PBS, resuspended in RLT buffer and stored at −80°C until nucleic acid extraction.

## Bioimaging

Cells were pelleted, washed in PBS, and fixed in 4% paraformaldehyde (Sigma-Aldrich) for 1 hour at room temperature. Cells were permeabilised using 0.5% Triton X-100 (Sigma-Aldrich) for 30 minutes at room temperature, blocked with 4% bovine serum albumin (Sigma-Aldrich) for 30 minutes at room temperature on a rotating platform and then labelled with a primary mouse monoclonal antibody against the IBDV VP2 protein (clone JF7-PD5) and incubated for 1 hour at room temperature. After washing in PBS, the cells were incubated for 1 hour with a goat-anti-mouse secondary monoclonal antibody conjugated to Alexa Fluor® 488 (Thermo Fisher Scientific). Cells were counterstained with DAPI (Sigma-Aldrich) and adhered to glass cover slips (TAAB, Aldermaston, UK) that had been coated with CellTak (Fisher Scientific) by centrifugation. Cover slips were mounted onto glass slides and stained cells were viewed with a Leica SP5 confocal microscope.

## Nucleic acid extraction

Total RNA was extracted from mock-infected and IBDV-infected B cells using an RNeasy kit (Qiagen) according to the manufacturer’s instructions. On-column DNA digestion was performed using RNase-free DNase (Qiagen) to remove contaminating genomic DNA. RNA samples were quantified using a Nanodrop Spectrophotometer (Thermo Scientific) and checked for quality using a 2100 Bioanalyzer (Agilent Technologies). All RNA samples had an RNA integrity number (RIN) ≥ 9.6. RNA samples were halved and processed for either microarrays or RTqPCR.

## Quantification of virus replication by RTqPCR

Reverse transcription of RNA samples was performed using SuperScript™ III Reverse Transcriptase (Thermo Fisher Scientific). The cDNA was synthesised according to the manufacturer’s instructions. Forward and reverse primers and a Taqman probe (Table S1) targeting a conserved region of the IBDV VP4 sequence (Sigma-Aldrich) were used to amplify the template. Briefly, the template, primers and probe were added to a Taqman Universal PCR Master Mix (Applied Biosystems) and reactions were performed on 7500 Fast Real-Time PCR system (Life Technologies) using the following cycling conditions: 95°C for 10 minutes; 40 cycles of 95°C for 15 seconds, 60°C for 1 minute; 95°C for 15 seconds; 60°C for 1 minute; 95°C for 30 seconds; 60°C for 15 seconds. All target gene expression levels were normalised to the housekeeping gene TBP and compared with the mock controls using the comparative C_T_ method (also referred to as the 2^−ΔΔCT^ method) (51).

## Microarrays

For the microarray study the GeneChip® 3’ IVT Express Kit (Affymetrix) was used following the manufacturer’s instructions. Hybridisation of RNA to chips and scanning of arrays was performed by the Medical Research Council’s Clinical Sciences Centre (CSC) Genomics Laboratory, Hammersmith Hospital, London, UK. RNA was hybridised to GeneChip Chicken Genome Array chips (Affymetrix) in a GeneChip Hybridization Oven (Affymetrix), the chips were stained and washed on a GeneChip Fluidics Station 450 (Affymetrix), and the arrays were scanned in a GeneChip Scanner 3000 7G with autoloader (Affymetrix).

## Nucleic acid labelling

Biotinylated fragmented RNA was prepared for each sample using standard procedures in GeneChip ® 3 IVT Express Kit User’s manual.

## Nucleic acid hybridization to array

Array hybridisation was performed according to the manufacturer’s instructions (Affymetrix). Labelled samples were hybridized to the GeneChip Chicken Genome Arrays (Affymetrix Inc.) in a GeneChip Hybridization Oven for 16 h at 45°C and 60 rpm in an Affymetrix Hybridization Oven 645.

## Array scanning and feature extraction protocol

After washing and staining, the arrays were scanned with the Affymetrix GeneChip Scanner 3000 7G. Gene-level expression signal estimates were derived from CEL files generated from raw data using the multi-array analysis (RMA) algorithm implemented from the Affymetrix GeneChip Command Console Software Version 3.0.1.

## Normalisation data transformation protocol

Data pre-processing and filtering was done using the Partek Genomics Suite software, v.6.6 and included: RMA background correction, quantile normalisation across all chips in the experiment, log_2_ transformation and median polish summarisation.

## Microarray data analysis

A one-way ANOVA (variable: treatment) adjusted with the Benjamini–Hochberg multiple-testing correction (false discovery rate (FDR) of *P*<0.05) was performed with Partek Genomics Suite (v6.6, Partek) across all samples. Principal component analysis (PCA) which shows sample variability in three-dimensional space confirmed lack of variability within infected samples (Fig. S1). Comparisons were conducted between B-cells infected with D78 and UK661 versus mock-treated cells and between cells infected with D78 versus cells infected with UK661. The analysis cut off criteria were fold change ≥ ± 1.5 and *P*-value ≤ 0.05. The Affymetrix chicken genome arrays contain probe sets for detecting transcripts from 17 avian viruses, including IBDV, allowing confirmation of viral infection. The full results of the microarray analysis are listed in Tables S2-S4. According to the MIAME guidelines, the original microarray data produced in this study have been deposited in the public database ArrayExpress (<u>http://www.ebi.ac.uk/microarray-as/ae/</u>), with the accession number: E-MTAB-5947.

Data mining and enrichment analysis was performed using the MetaCore software suite (Clarivate Analytics, <u>https://clarivate.com/products/metacore/</u>). Enrichment analysis consisted of mapping gene IDs of the datasets onto gene IDs of human orthologues in entities of built-in functional ontologies represented in MetaCore by pathway maps and process networks. Statistical significance was measured by the number of genes that map onto a given pathway and was calculated on the basis of p-value, based on hypergeometric distribution (a built-in feature of MetaCore). Enrichment analysis for the mock-infected Vs D78, mock-infected Vs UK661 and D78 Vs UK661 datasets is presented in Fig. S2-S4.

## Microarray validation

RTqPCR was performed on RNA samples using a two-step procedure. RNA was first reverse-transcribed into cDNA using the QuantiTect Reverse Transcription Kit (Qiagen) according to the manufacturer’s instructions. qPCR was then conducted on the cDNA in a 384-well plate with a ABI-7900HT Fast qPCR system (Applied Biosystems). Mesa Green qPCR MasterMix (Eurogentec) was added to the cDNA (5μl for every 2μl of cDNA). The following amplification conditions were used: 95°C for 5 minutes; 40 cycles of 95°C for 15 seconds, 57°C for 20 seconds, and 72°C for 20 seconds; 95°C for 15 seconds; 60°C for 15 seconds; and 95°C for 15 seconds. Primer sequences for genes that were used in the study are given in Supplementary Table 1. The output Ct values and dissociation curves were analysed using SDS v2.3 and RQ Manager v1.2 (Applied Biosystems). Gene expression data were normalised against the housekeeping gene GAPDH, and compared with the mock controls using the comparative C_T_ method (51). All samples were loaded in triplicate.

## Acknowledgements

We are grateful for the skilled support of Laurence Game and Ivan Andrew of the Medical Research Council’s (MRC) Clinical Sciences Centre’s (CSC) Genomics Laboratory in conducting microarray analysis. In addition, we thank Nicolas Eterradossi of ANSES France for supplying IBDV strains, and for useful discussions.

## Ethical statement

All animal procedures were performed in accordance with the UK Animal (Scientific Procedures) Act 1986 under Home Office Establishment, Personal and Project licences, after the approval of the Animal Welfare and Ethical Review Body (AWERB) at The Pirbright Institute.

---

**Supplementary Figure 1.**
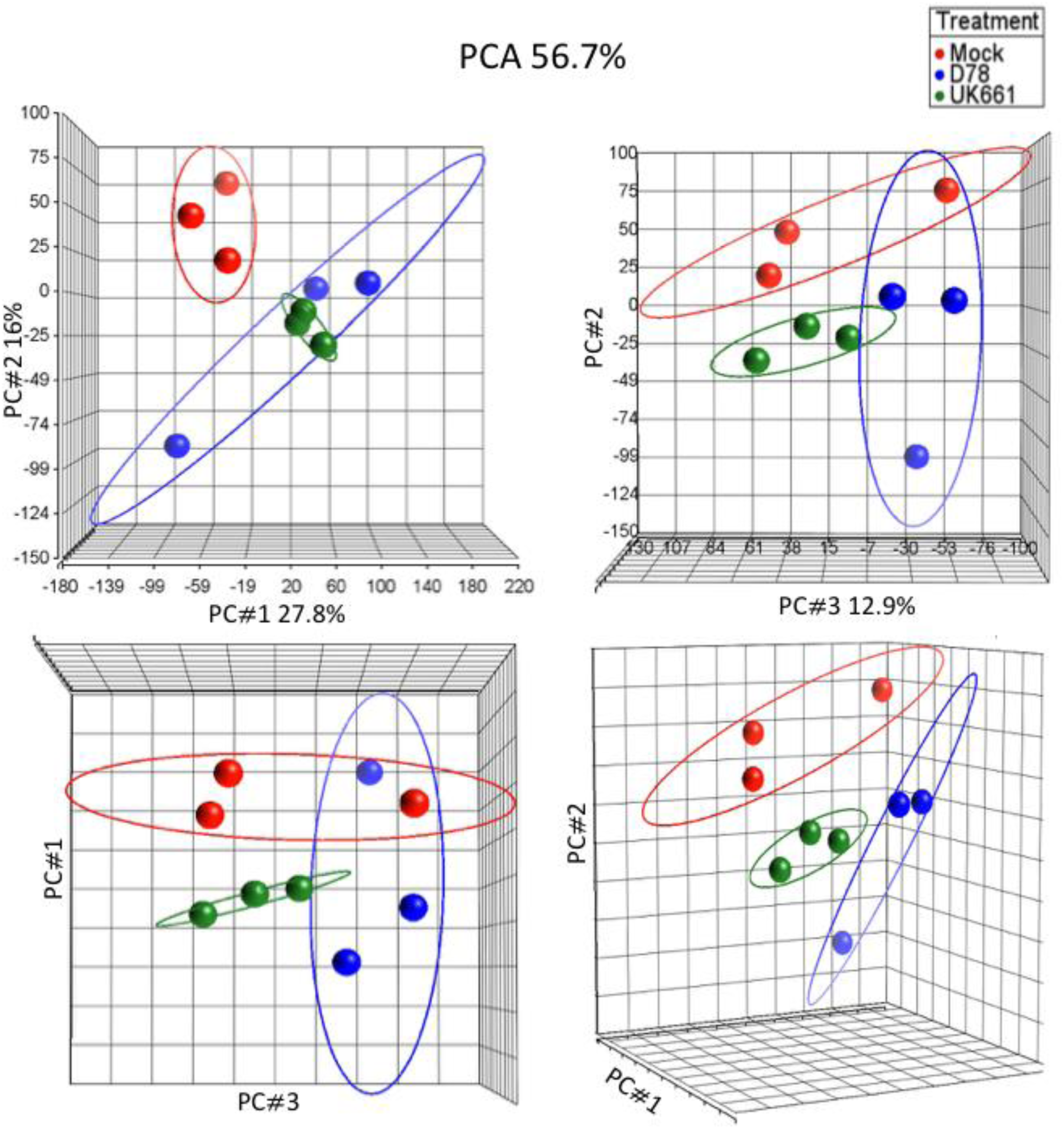
Four alternative views of three-dimensional scatter PCA plots displaying the similarity of expression of samples as indicated by dots. Bounding ellipses and colours represent mock-(red), D78-(blue) and UK661-(green) infected samples.

**Supplementary Figure 2.**
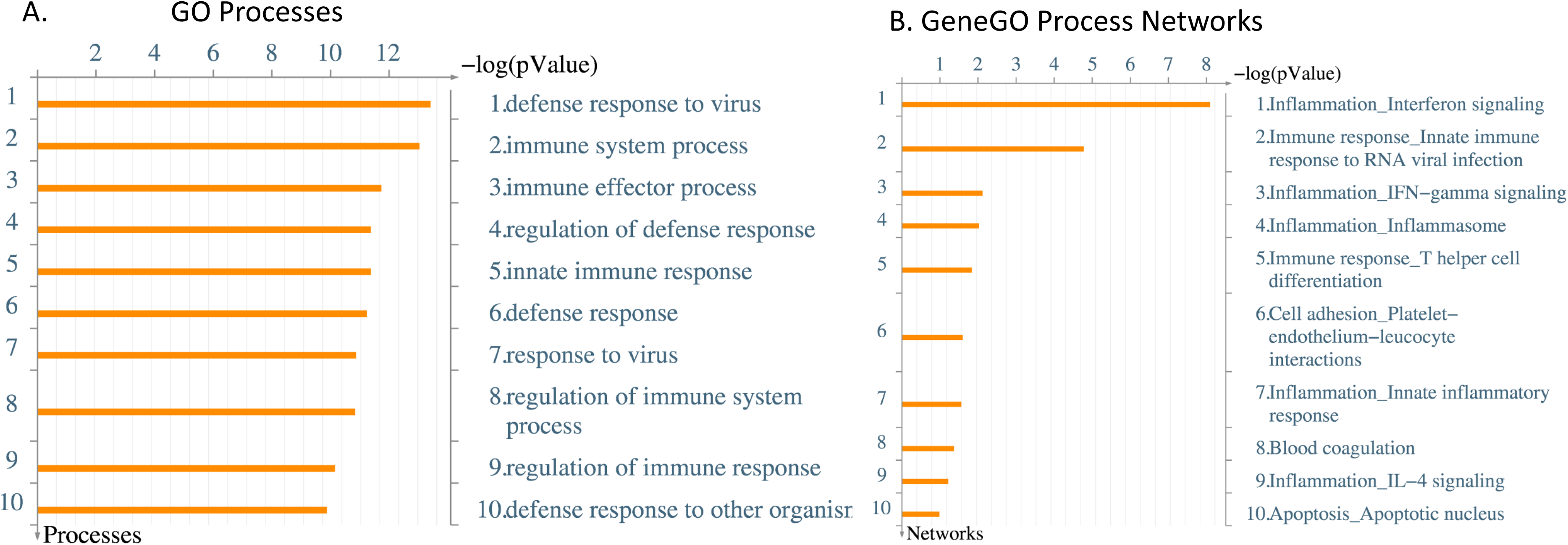
Enrichment analysis of the genes identified in the microarray comparison D78 Vs Mock by MetaCore: (A) GO Processes, (B) Process networks. The p-value is estimated by MetaCore ‘Statistically significant’ method. MetaCore version 6.31 build 68930.

**Supplementary Figure 3.**
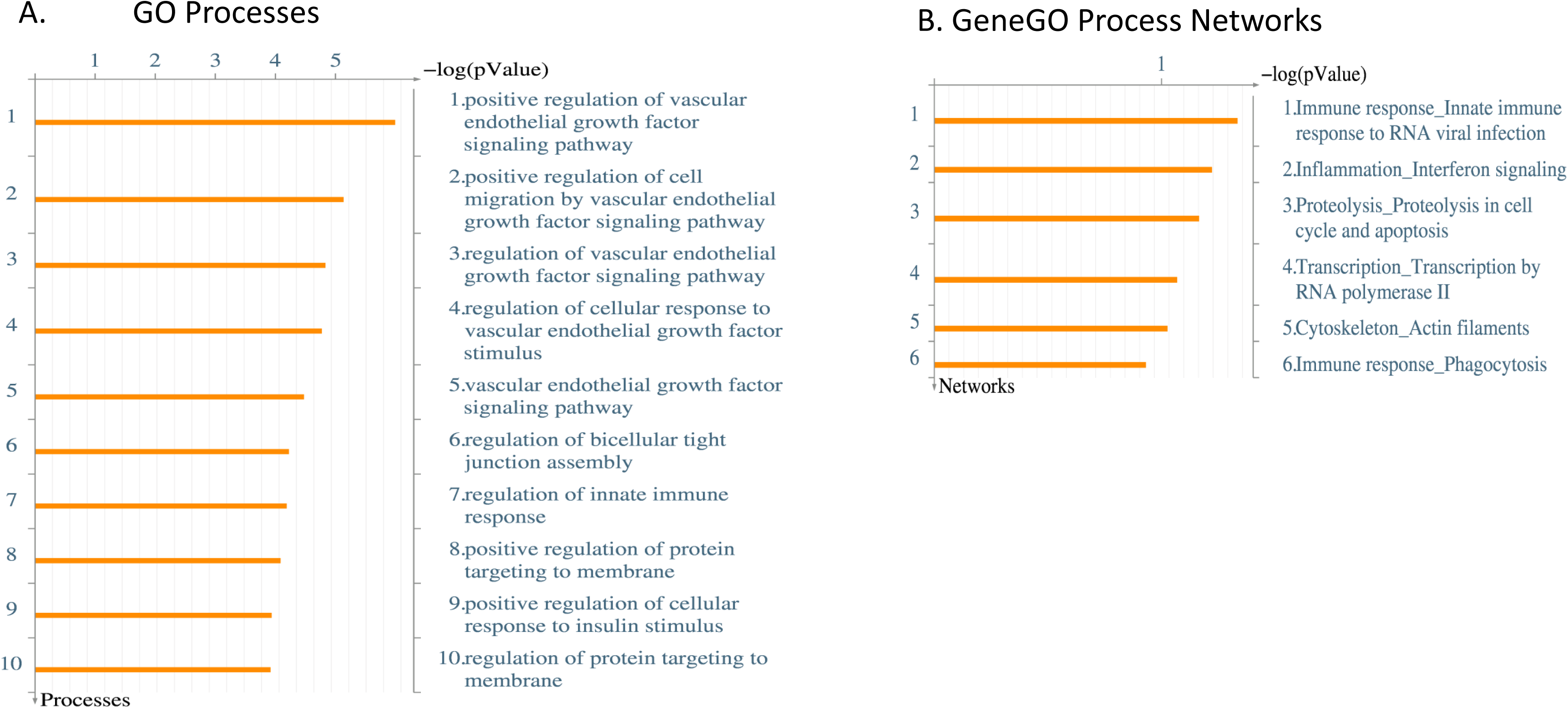
Enrichment analysis of the genes identified in the microarray comparison UK661 Vs Mock by MetaCore: (A) GO Processes, (B) Processes networks. The p-value is estimated by MetaCore ‘Statistically significant’ method. MetaCore version 6.31 build 68930.

**Supplementary Figure 4.**
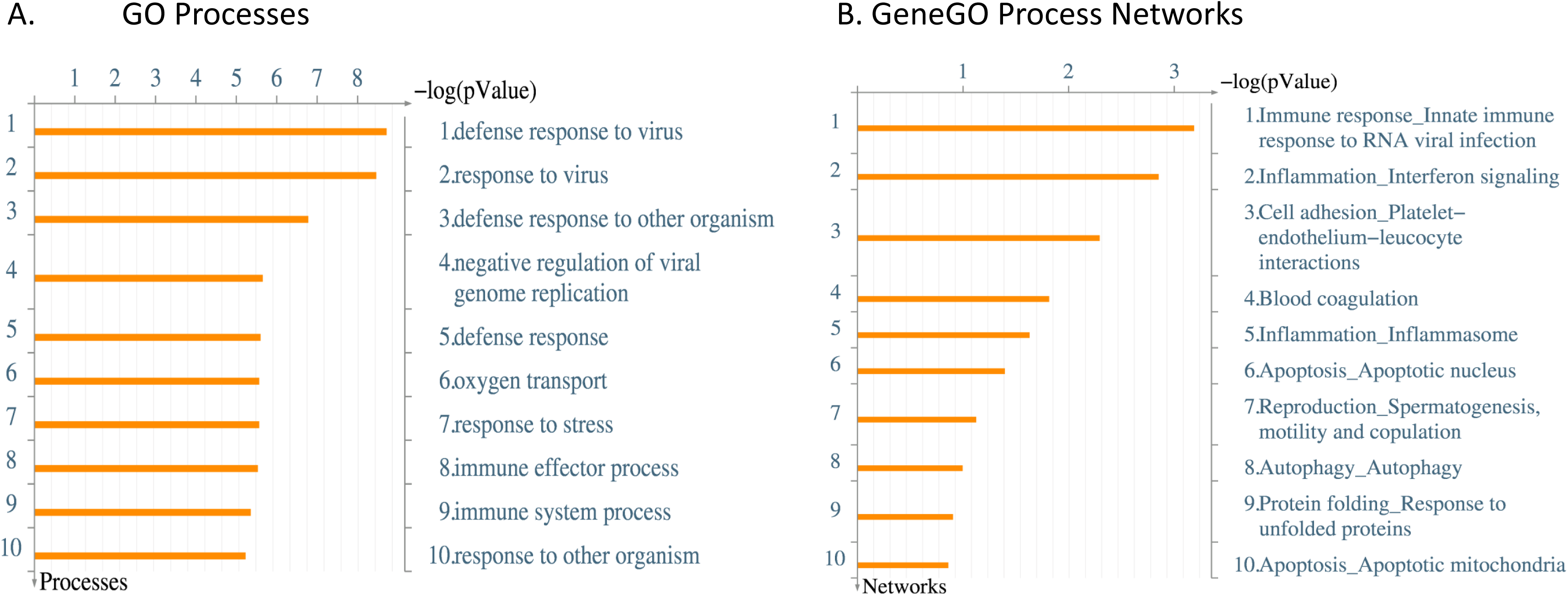
Enrichment analysis of the genes identified in the microarray comparison D78 Vs UK661 by MetaCore: (A) GO Processes, (B) Process networks. The p-value is estimated by MetaCore ‘Statistically significant’ method. MetaCore version 6.31 build 68930

**Supplementary Table 1.**
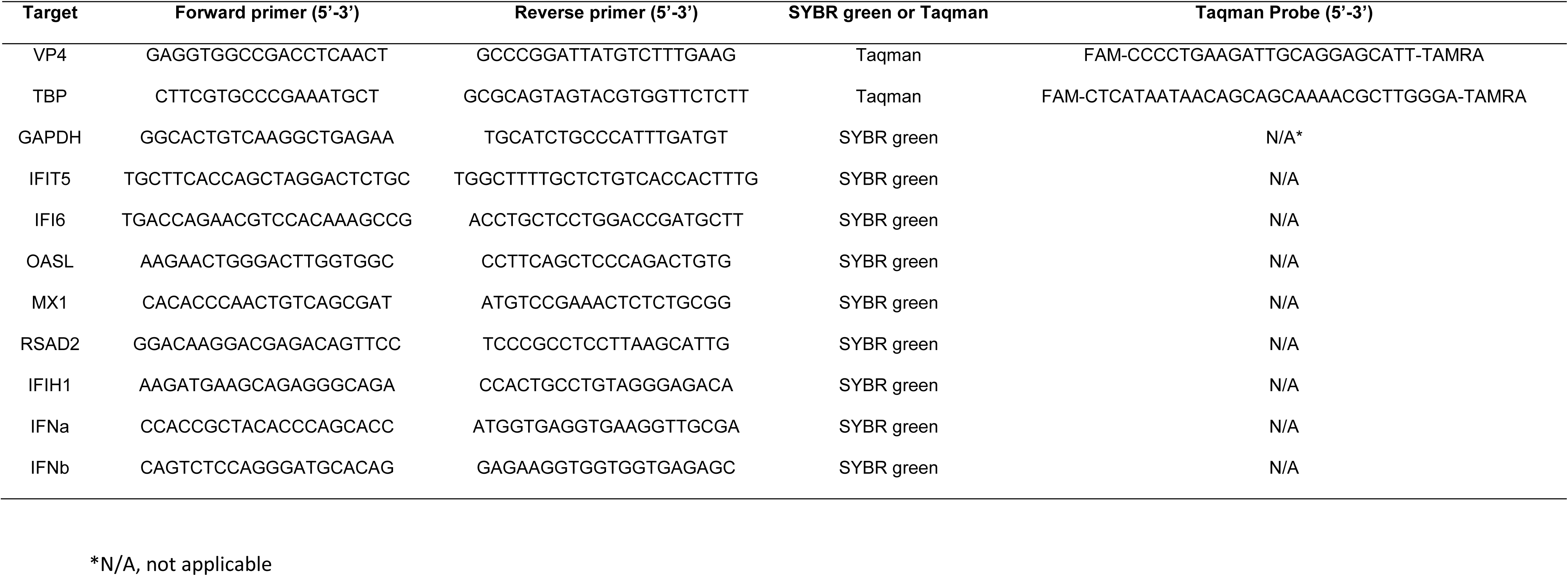
Primer and probe sequences used in quantitative PCR.

**Supplementary Table 2.**
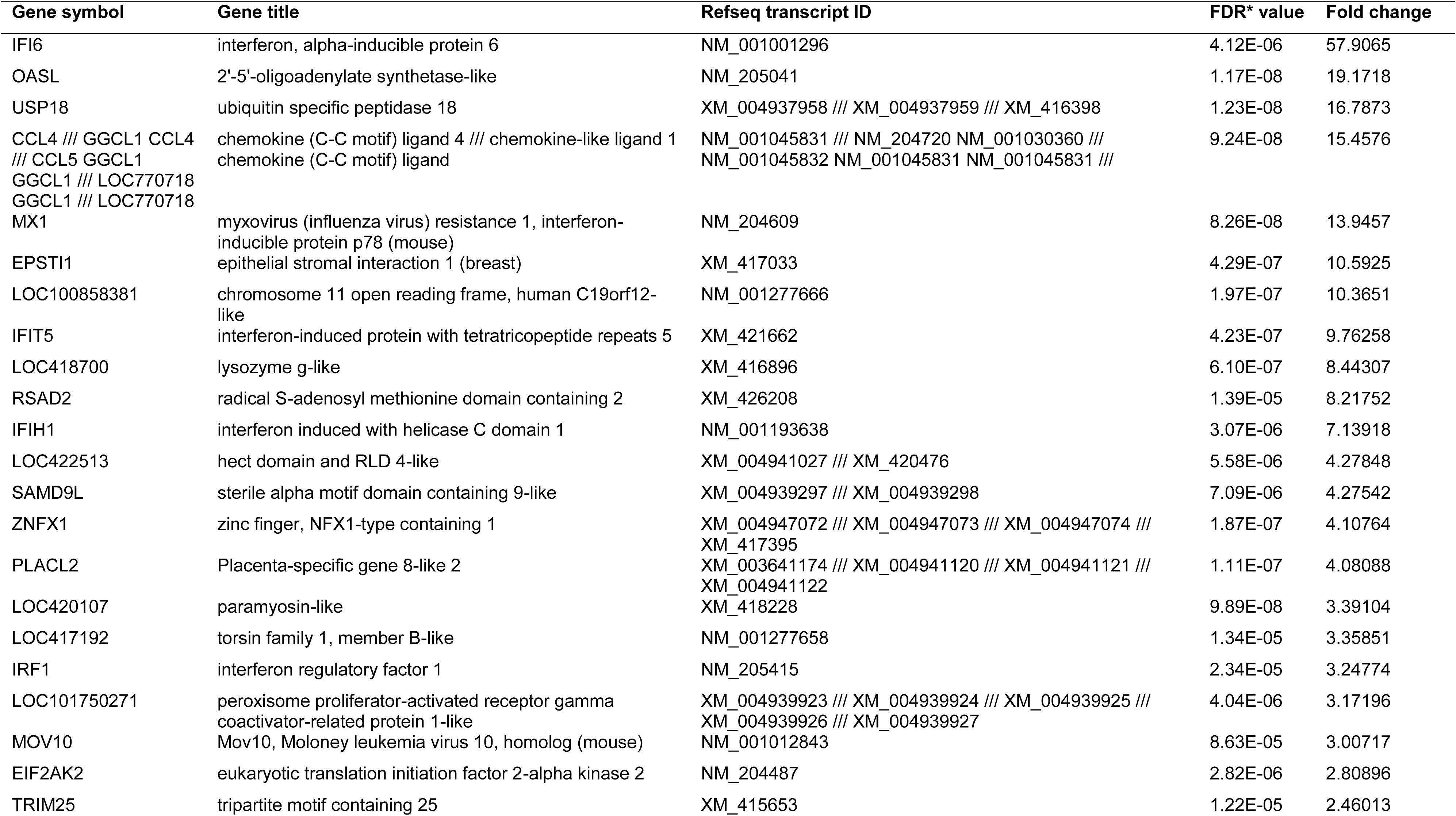
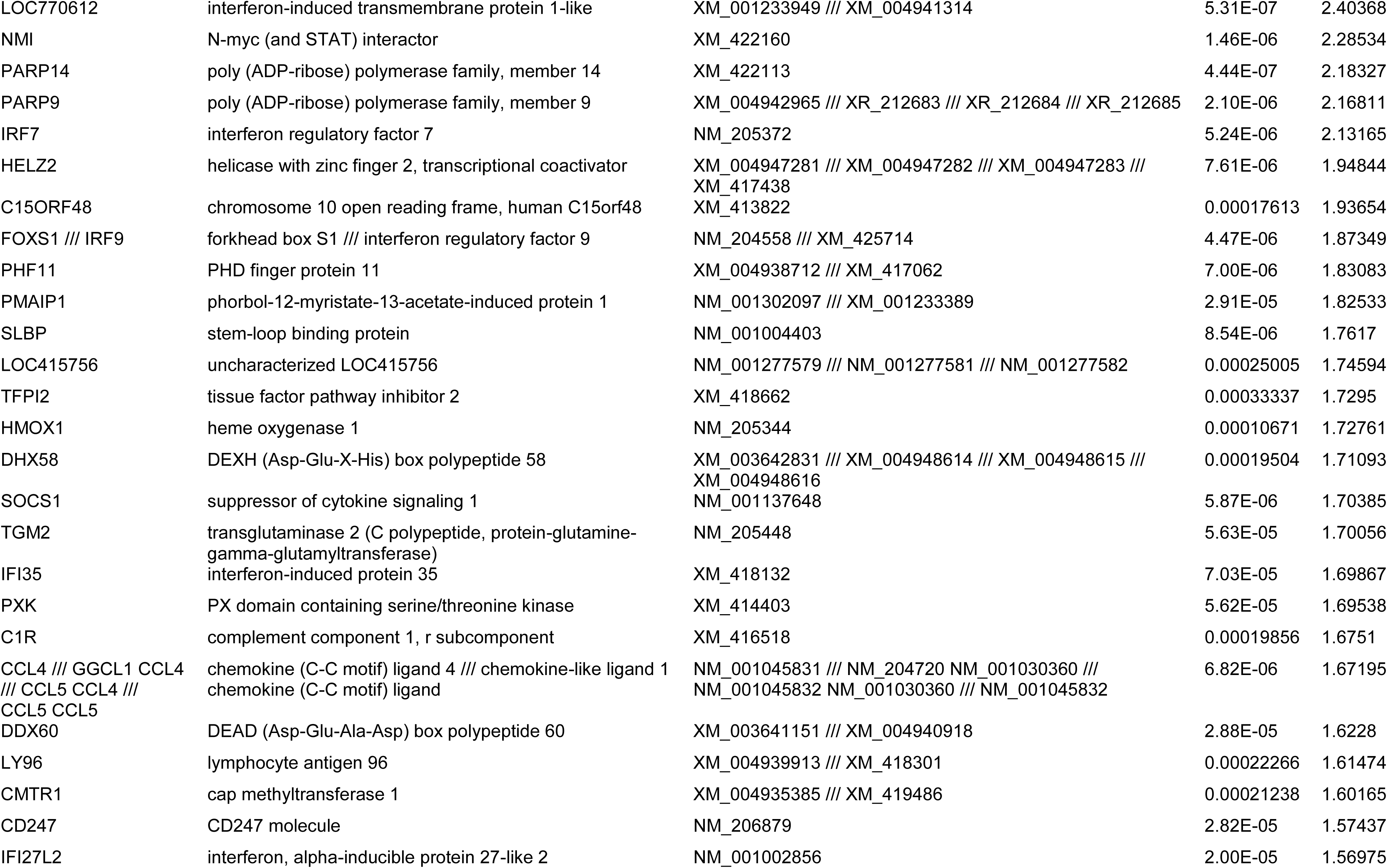
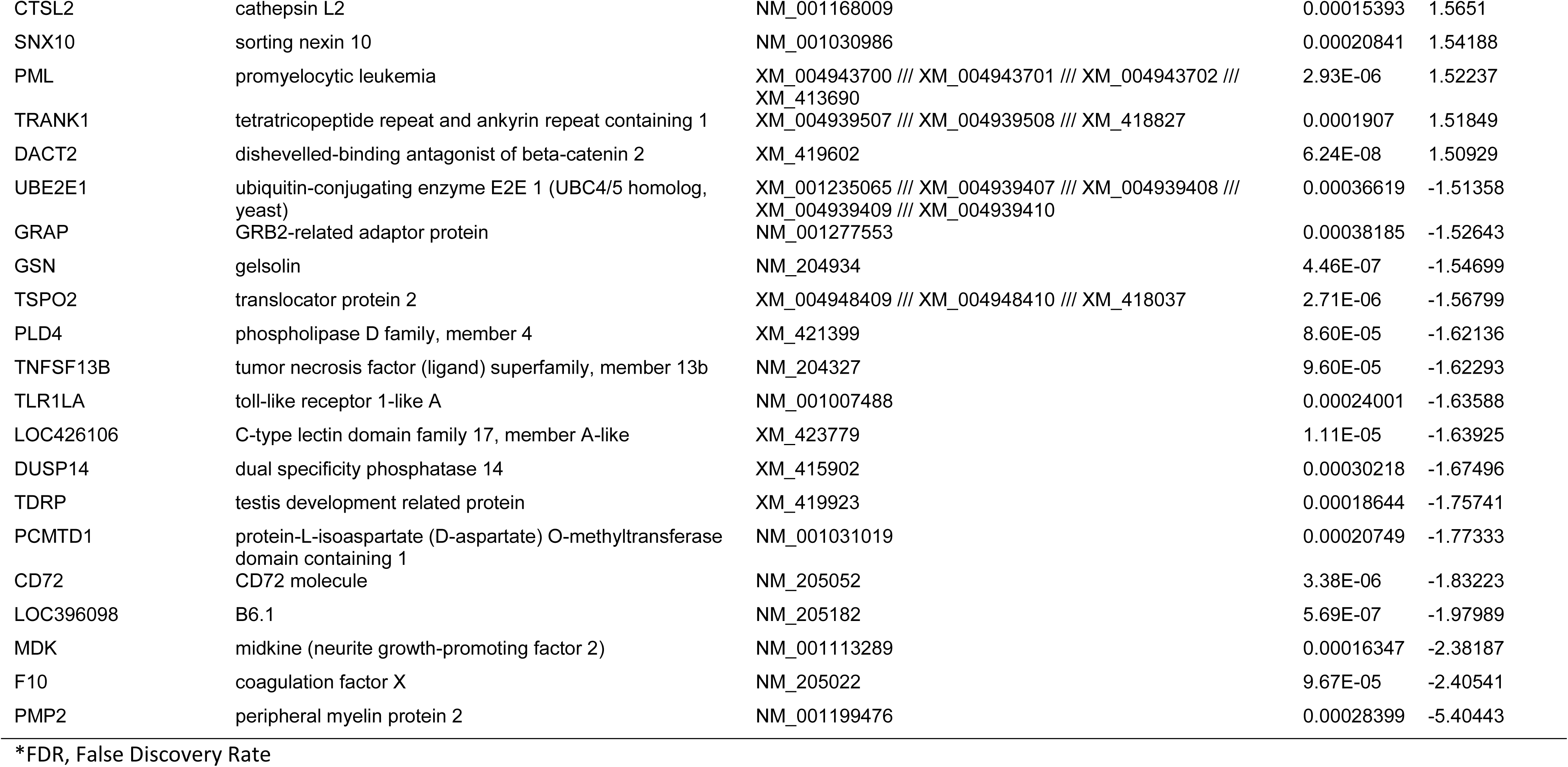
Microarray data, D78 Vs Mock.

**Supplementary Table 3.**
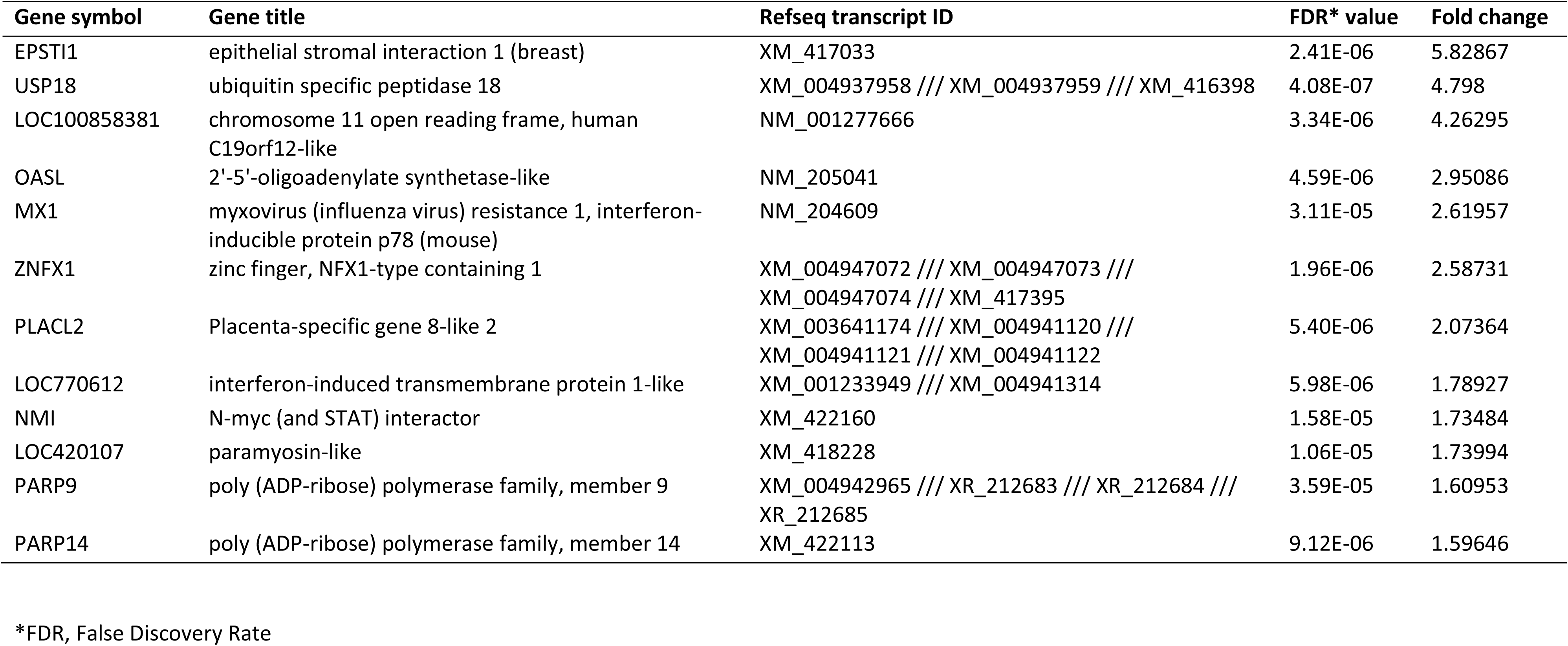
Microarray data, UK661 Vs Mock.

| Gene symbol | Gene title | Refseq transcript ID | FDR* value | Fold change |
| --- | --- | --- | --- | --- |
| EPSTI1 | epithelial stromal interaction 1 (breast) | XM_417033 | 2.41E-06 | 5.82867 |
| USP18 | ubiquitin specific peptidase 18 | XM_004937958 /// XM_004937959 /// XM_416398 | 4.08E-07 | 4.798 |
| LOC100858381 | chromosome 11 open reading frame, human<br>C19orf12-like | NM_001277666 | 3.34E-06 | 4.26295 |
| OASL | 2'-5'-oligoadenylate synthetase-like | NM_205041 | 4.59E-06 | 2.95086 |
| MX1 | myxovirus (influenza virus) resistance 1, interferon-inducible protein p78 (mouse) | NM_204609 | 3.11E-05 | 2.61957 |
| ZNFX1 | zinc finger, NFX1-type containing 1 | XM_004947072 /// XM_004947073 ///<br>XM_004947074 /// XM_417395 | 1.96E-06 | 2.58731 |
| PLACL2 | Placenta-specific gene 8-like 2 | XM_003641174 /// XM_004941120 ///<br>XM_004941121 /// XM_004941122 | 5.40E-06 | 2.07364 |
| LOC770612 | interferon-induced transmembrane protein 1-like | XM_001233949 /// XM_004941314 | 5.98E-06 | 1.78927 |
| NMI | N-myc (and STAT) interactor | XM_422160 | 1.58E-05 | 1.73484 |
| LOC420107 | paramyosin-like | XM_418228 | 1.06E-05 | 1.73994 |
| PARP9 | poly (ADP-ribose) polymerase family, member 9 | XM_004942965 /// XR_212683 /// XR_212684 ///<br>XR_212685 | 3.59E-05 | 1.60953 |
| PARP14 | poly (ADP-ribose) polymerase family, member 14 | XM_422113 | 9.12E-06 | 1.59646 |

\*FDR, False Discovery Rate

**Supplementary Table 4.**
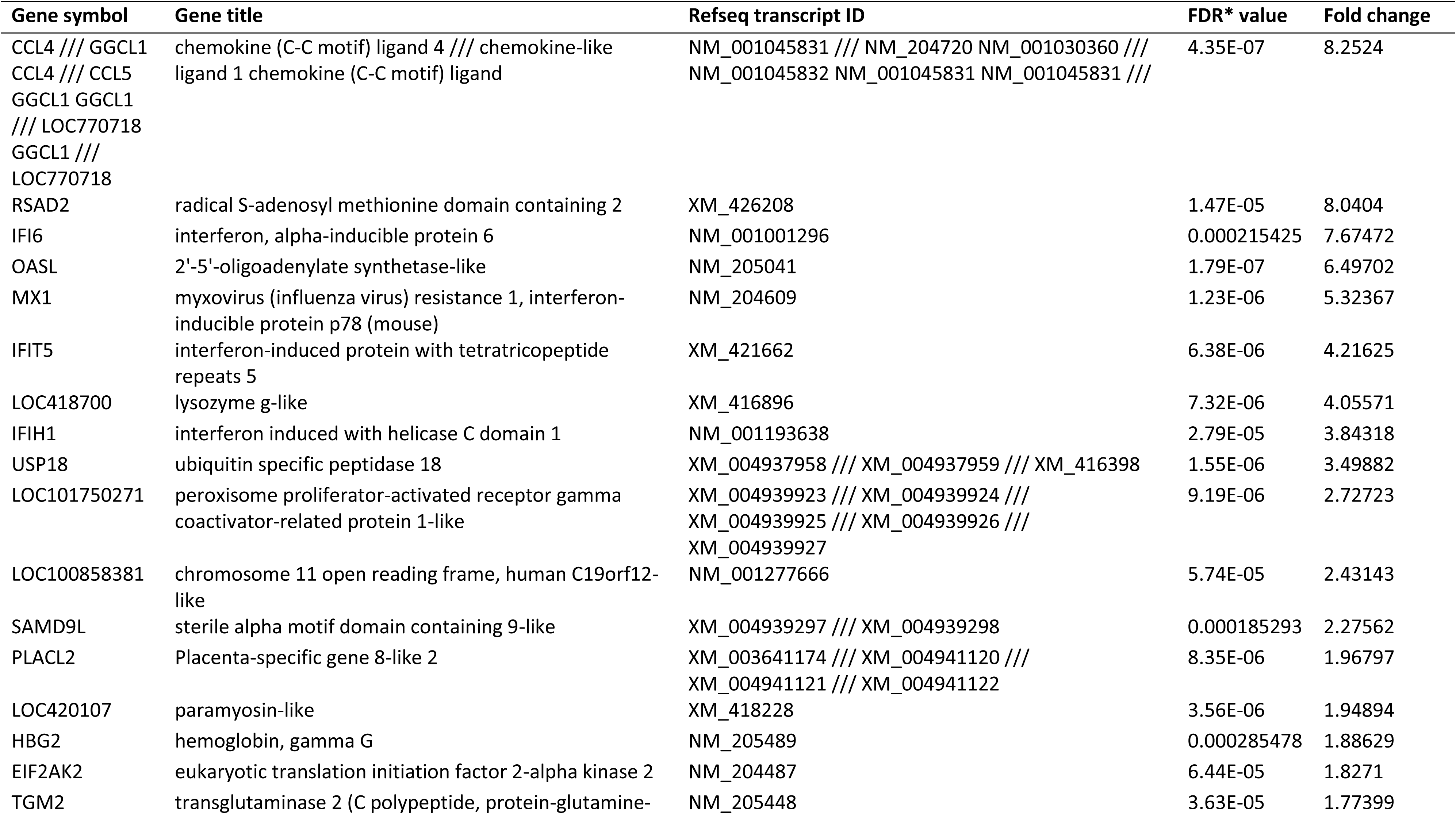
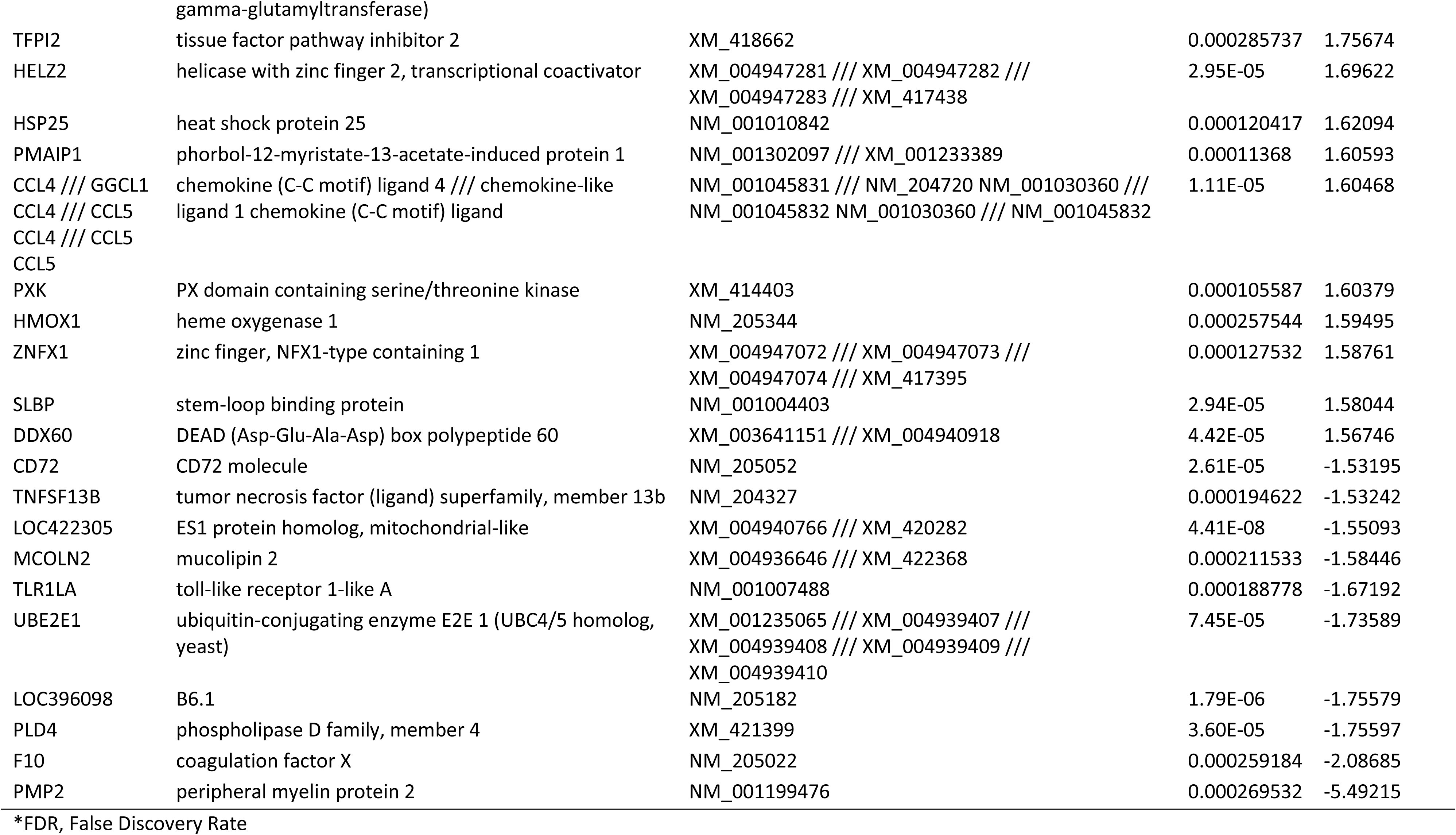
Microarray data, D78 Vs UK661.

